# Identification of teleost *tnnc1a* enhancers for specific pan-cardiac transcription

**DOI:** 10.1101/2024.02.26.582099

**Authors:** Jianhui Bai, Xiangyun Wei

## Abstract

Troponin C regulates muscle contraction by forming the troponin complex with troponin I and troponin T. Different muscle types express different troponin C genes. The mechanisms of such differential transcription are not fully understood. The Zebrafish tnnc1a gene is restrictively expressed in cardiac muscles. We here identify the enhancers and promoters of the zebrafish and medaka *tnnc1a* genes, including intronic enhancers in zebrafish and medaka and an upstream enhancer in the medaka. The intronic and upstream enhancers are likely functionally redundant. The GFP transgenic reporter driven by these enhancers is expressed more strongly in the ventricle than in the atrium, recapitulating the expression pattern of the endogenous zebrafish *tnnc1a* gene. Our study identifies a new set of enhancers for cardiac-specific transgenic expression in zebrafish. These enhancers can serve as tools for future identification of transcription factor networks that drive cardiac-specific gene transcription.

## Introduction

Troponin C (TnC) regulates muscle contraction by forming the troponin complex with troponin I and troponin T to modulate the cross-bridge interactions between myosin and actin filaments. This modulation is mediated by a series of conformational changes in the troponin complex, which are initiated by troponin C’s sensing and binding to calcium ions (Fig. 1A) (Farah and Reinach, 1995; Gomes et al., 2002) (Vinogradova et al., 2005). In vertebrates, multiple troponin C genes exist and are differentially expressed in striated muscles. In mammals, cardiac and slow-twitch skeletal muscles express the *tnnc1* gene (Gahlmann et al., 1988), which codes for cardiac troponin C (cTnC), whereas fast-twitch skeletal muscles express the *tnnc2* gene, which codes for fast-twitch skeletal troponin C (fsTnC) (Fig. 1B) (Berezowsky and Bag, 1988; Schreier et al., 1990). In teleost, fast-twitch skeletal muscles express *tnnc2* as in mammals. However, the situation for cardiac and slow-twitch muscles is more complex because there are two *tnnc1* genes due to the teleost-specific genome duplications during evolution (Genge et al., 2016). *tnnc1a* is restrictively expressed in cardiac muscles, and *tnnc1b* is expressed in both the cardiac and slow-twitch skeletal muscles (Genge et al., 2013; Sogah et al., 2010). The myocardial expressions of *tnnc1a* and *tnnc1b* also vary between the ventricular and atrial heart chambers depending on fish species and thermal acclimation (Genge et al., 2013) (Fig. 1B).

**Figure 1.**
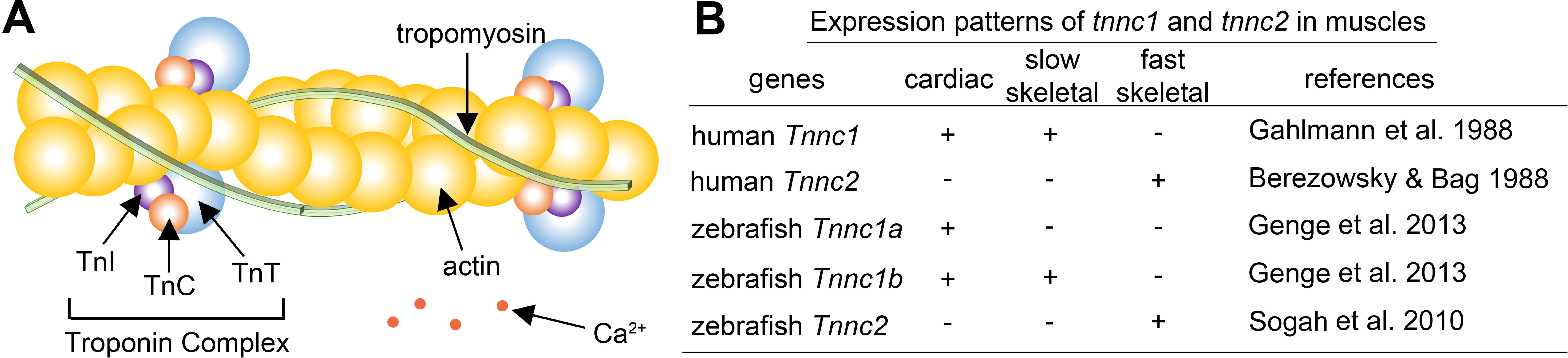
The function and expression patterns of TnC in vertebrates. A. A schematic of the Troponin complex in regulating muscle contraction. The binding of Ca2+ to TnC leads to a series of conformational changes that eventually translocate tropomyosin from the outer domain of actin filaments, enabling the binding and sliding of myosin (not shown) along actin filaments to generate contraction. B. A table illustrates the differential expression of human and zebrafish tnnc1 and tnnc2 among striated muscles.

Because different troponin C proteins have different sensitivities to Ca^2+^ (Churcott et al., 1994; Harrison and Bers, 1989), these differential expression patterns of troponin C genes contribute directly to the differences in contractile properties among the three types of striated muscles as well as between cardiac ventricle and atrium (Chien et al., 1993). Thus, transcriptional regulation of the cardiac expression of *tnnc1*, *tnnc1a*, and *tnnc1b* is an essential aspect of the biology of cardiac development and function (Tabibiazar et al., 2003).

Some progress has been made in understanding the transcriptional regulation of mammalian *tnnc1*. An upstream enhancer and an intronic enhancer were identified for the expression of *tnnc1* in cardiac and slow-twitch skeletal muscles, respectively (Christensen et al., 1993; Parmacek et al., 1994; Parmacek et al., 1992; Schreier et al., 1990). These enhancers interacted directly or indirectly in cell culture with transcription factors (TFs) MyoD, SP1, GATA4, and HMG2 to regulate reporter gene expression (Christensen et al., 1993; Ip et al., 1994; Montgomery and Litvin, 1997; Parmacek et al., 1994). However, it is unclear whether *tnnc1*’s cardiac enhancer interacts with other key cardiac TFs, such as Nkx2.5, Hand2, and MEF2 (Stainier, 2001; Staudt and Stainier, 2012; Yelon, 2001). Thus, how cis- and trans-regulatory elements regulate mammalian TnC expression remains to be fully understood.

A better understanding of TnC expression can be facilitated by unraveling and comparing the transcriptional regulation of TnC of various animal species via phylogenetic and comparative genomic approaches. To this end, teleost *tnnc1a* and *tnnc1b* genes are good models to study because the greater diversity of teleost provides a rich source for such investigation (Glasauer and Neuhauss, 2014). Thus, we investigated the cis-regulation of teleost *tnnc1a* in this study. Here, we report the identification of the promoters and enhancers of medaka and zebrafish *tnnc1a* genes. In addition, the conserved motifs of *tnnc1a* enhancers were assessed by mutation analysis and compared with those of enhancers of 22 other cardiac genes. Our study not only identified new pan-cardiac enhancers useful for transgenic research but also gained insights into the general principles underlying pan-cardiac transcription.

## Materials and Methods

### Animal care and maintenance

Tubingen (TU) wildtype zebrafish, transgenic zebrafish, and medaka (obtained from a local pet store) were raised and maintained on a 14 h light/10 h dark cycle. Zebrafish embryos were raised at 28.5 °C in an incubator. In this study, both male and female fish were used for experiments, although the sexes were not distinguishable in zebrafish larvae. Animal care and handling were in accordance with the guidelines of the University of Pittsburgh

### In situ hybridization analysis of the mRNA expression of the *tnnc1a* gene

*In situ* hybridization was processed according to previously-published standard protocol (Thisse and Thisse, 2008; Zou et al., 2006), with the following specific adjustments: Zebrafish embryos were first treated with 0.003% phenylthiourea (PTU) between 1 and 3 dpf (days postfertilization) to block melanin pigmentation. 30-, 48-, and 72-hpf (hours postfertilization) zebrafish embryos were treated with 1ug/ml proteinase K for 30 min. To make specific riboprobes against *tnnc1a*, a cDNA library was first made by reverse transcription using the SuperScript® III First-Strand Synthesis System Kit (Thermofisher, catalog #: 18080051), random hexamer primers, and RNA purified from 5-dpf embryos with the miRNeasy Mini Kit (Qiagen, catalog #: 217004). Using this cDNA library, a 406-bp fragment of the 486-bp coding sequence (CDS) of *tnnc1a* (GenBank: AF434188.1) was amplified by PCR using *tnnc1a*-CDS primers (Table 1) and cloned into the pGEM®-T Easy Vector Systems (Promega, catalog #: A1360). A *tnnc1a* cDNA clone was confirmed by sequencing and used to make digoxigenin-labeled riboprobes with the Ambion™ MAXIscript™ SP6/T7 In Vitro Transcription Kit (Thermofisher, catalog #: AM1322) and a digoxigenin labeling mix (Roche, catalog#: 11277073910). The synthesized riboprobes were treated with DNase I digestion to remove the DNA template, purified with an RNeasy mini kit (Qiagen, catalog #: 74104), and dissolved in 50% formamide for storage at -80°C.

NBT/BCIP chromogenic staining of embryos was performed at 4℃ for 24-72 hours.

Images were taken with a Qimaging Retiga 1300 camera under a Leica M205FA fluorescence microscope.

### Identification of candidates for cis-regulatory elements

Candidate cis-regulatory elements of the *tnnc1a* genes were identified among conserved non-coding sequences using the method described previously (Fang et al., 2020). Specifically, the conserved noncoding sequences in teleost *tnnc1a* genes were revealed by the PhastCons histograms of Multiz alignments of five fish species at the UCSC genome browser: medaka (*Oryzias latipes*), zebrafish (*Danio rerio*), stickleback (*Gasterosteus aculeatus*), fugu (*Takifugu rubripes*), tetraodon (*Tetraodon nigroviridis*) on the UCSC BLAT search genome browser (https://genome.ucsc.edu). The basal promoter (also known as core promoter) is expected to reside in the conserved noncoding regions within 100 bp from the transcription start site, whereas the enhancers are expected to reside in conserved noncoding regions in distance, particularly in a single stretch of 80–250 bp, either upstream or downstream of the transcription start site.

### Generation of transgene constructs

To test the transcriptional activities of DNA fragments amplified by PCR (Table 1), they were cloned between the Apa I and Asis I sites of an I-SceI meganuclease-based enhancer-testing transgenic GFP reporter vector (Fang et al., 2017), which contains the elements of the reporter gene in the following 5’ to 3’ order: a Kpn I site, either the basal promoter region of the *ponli* gene as published previously (Fang et al., 2017) or the basal promoter region of the zebrafish *tnnc1a* gene (from -109 bp upstream of the transcription start site to 160 bp downstream of the 5’ splicing site of *tnnc1a* intron 1, which includes the 56-bp 5’UTR and 58-bp coding sequence of *tnnc1a* exon 1), the Apa I and Asis I sites, the 474-bp 3’ end of *ponli* intron 1), the noncoding region of *ponli* exon 2 (67 bp), a Kozak sequence (GCCACC), GFP reporter CDS, and a 3’-UTR with an sv40 polyadenylation signal. Translation can be initiated at the start codon after the partial *ponli* exon 2 and the Kozak sequence.

### *In vivo* transgenic assay of enhancer candidates and generation of stable transgenic fish

The transcriptional activities of transgenic reporter constructs were tested in zebrafish embryos by embryonic injection. The injection solution comprised 1μl of plasmid, 1μl of ISce1 restriction enzyme buffer, 1μl of I-Sce1, and 7μl of water. Zebrafish embryos were injected roughly with 20 pg of a transgene construct and 0.01 unit of I-SceI at the 1-cell stage. The GFP-expressing founder fish were outcrossed with wildtype fish to screen for stable transgenic fish. The GFP expression was examined under a Leica M205FA fluorescence microscope.

### Truncation and mutation analysis

The PCR-based Q5 Site-Directed Mutagenesis Kit (NEB, #E0554S) was used to perform truncation (constructs L-P) and substitution mutagenesis using primers specified in Table 1. The substitution sequences were unrelated sequences of the same length (12-16bp), containing either a SacI or KpnI site for verification by restriction enzyme digestion. The substitution sequences were verified not to contain known transcription factor binding sites by using TOMTOM (http://meme-suite.org/tools/tomtom). All constructs were verified by restriction enzyme digestion and sequencing.

### Identification of conserved sequence motifs

Conserved short sequence motifs were identified with web-based sequence comparison tools MEME (Bailey et al., 2009) and Improbizer. In using MEME (http://meme-suite.org), the range of motif length was set to be between 6 and 12 bp because most transcription factor binding sites are within this range. For Improbizer https://users.soe.ucsc.edu/~kent/improbizer/improbizer.html), the setting was adjusted to ignore location and to include reverse complementary sequences. Other parameters were at their default values. Putative transcription factor binding sites in the 509-bp zebrafish *tnnc1a* enhancer were predicted with Transfac (http://gene-regulation.com/pub/databases.html), verified by JASPAR (http://jaspar.genereg.net) using a relative profile score threshold of 80% or higher, and TOMTOM (http://meme-suite.org/tools/tomtom) (Bailey et al., 2009).

## Results

### *tnnc1a* is expressed in both the atrial and ventricular chambers of zebrafish embryonic heart

Previously, in situ analysis showed that *tnnc1a* was restrictively expressed in the ventricle of the zebrafish embryonic heart (Sogah et al., 2010). However, quantitative RT-PCR revealed that *tnnc1a* mRNA was expressed in both the ventricle and atrium of adult zebrafish heart (Genge et al., 2013). To verify the expression pattern of *tnnc1a* in zebrafish embryos, we performed in situ hybridization with a probe that recognizes 406 bp of the 486-bp CDS of *tnnc1a*, rather than the entire cDNA as used by Sogah et al (Sogah et al., 2010). We confirmed that *tnnc1a* is expressed exclusively in the heart in both the atrial and ventricular chambers (Fig. 2 A-C). At 30 hpf, when the atrioventricular canal starts to emerge to separate the S-shaped heart tube into the atrial and ventricular chambers (Peal et al., 2011) (Lombardo et al., 2019), the *tnnc1a* signals were observable in the entire heart tube (Fig. 2D, 2E). At 48 hpf, when the morphologically distinct ventricle is positioned more medially and the atrium is further left to the ventricle, *tnnc1a* was expressed in both the ventricle and atrium, with much stronger signals in the ventricle, a pattern very similar to zebrafish *mlc2* expression (Rottbauer et al., 2006). The much stronger ventricular signals made it easy to overlook atrial expression when in situ staining is not developed fully (Fig. 2D, E, F); this may explain why *tnnc1a’*s atrial expression in the embryonic heart was overlooked (Sogah et al., 2010). At 72 hpf, the atrial expression of *tnnc1a* remained, suggesting it is unlikely that the atrial signals were derived from residual mRNA expressed early on in the heart tube (Fig. 2H, 2I). Thus, as in the adult heart (Genge et al., 2013), zebrafish *tnnc1a* is expressed in both chambers of the embryonic heart, although much more abundantly in the ventricle.

**Figure 2.**
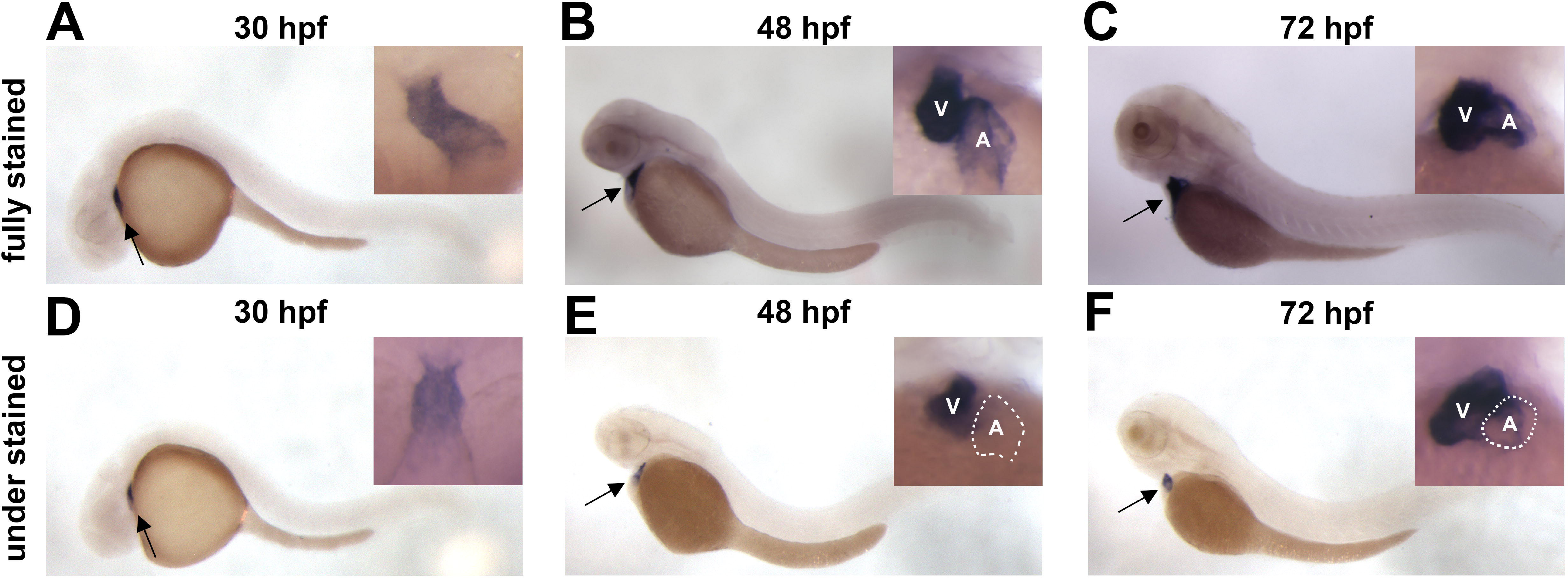
Cardiac expression of tnnc1a in embryonic zebrafish heart. **A-C.** Lateral views of fully stained in situ hybridization of *tnnc1a* mRNA in whole wildtype embryos at 30 hpf (A), 50 hpf (B), and 72 hpf (C). **D-F.** Underdeveloped in situ hybridization analysis, showing the lack of staining in the atrium. Insets show ventral views at higher magnifications. A, atrium; V, ventricle.

### Identification of teleost *tnnc1a* enhancer regions

Mammalian *tnnc1* is regulated by an upstream enhancer for its pan-cardiac expression and an intronic enhancer for its slow-twitch skeletal muscle expression (Christensen et al., 1993; Parmacek et al., 1994; Parmacek et al., 1992) (Fig. 1B). The restrictive pan-cardiac expression of zebrafish *tnnc1a* suggests that its enhancers may differ from those of mammalian *tnnc1*. To identify *tnnc1a*’s enhancers, we first searched for conserved non-coding sequences in the *tnnc1a* locus of zebrafish, medaka, stickleback, tetraodon, and fugu because cis-regulatory elements often reside in conserved non-coding regions (Kulaeva et al., 2012; Lenhard et al., 2012; Rouault et al., 2010). On the UCSC Genome Browser, we noticed highly conserved non-coding sequences in both the upstream region and intron 1 among medaka, stickleback, tetraodon, and fugu (Fig. 3). Of note, the sequence comparison at UCSC did not reveal corresponding sequence conservation at the zebrafish *tnnc1a* locus possibly because the zebrafish genome is too large and complex for zebrafish *tnnc1a* to be aligned with other fish’s *tnnc1a* genes.

**Figure 3.**
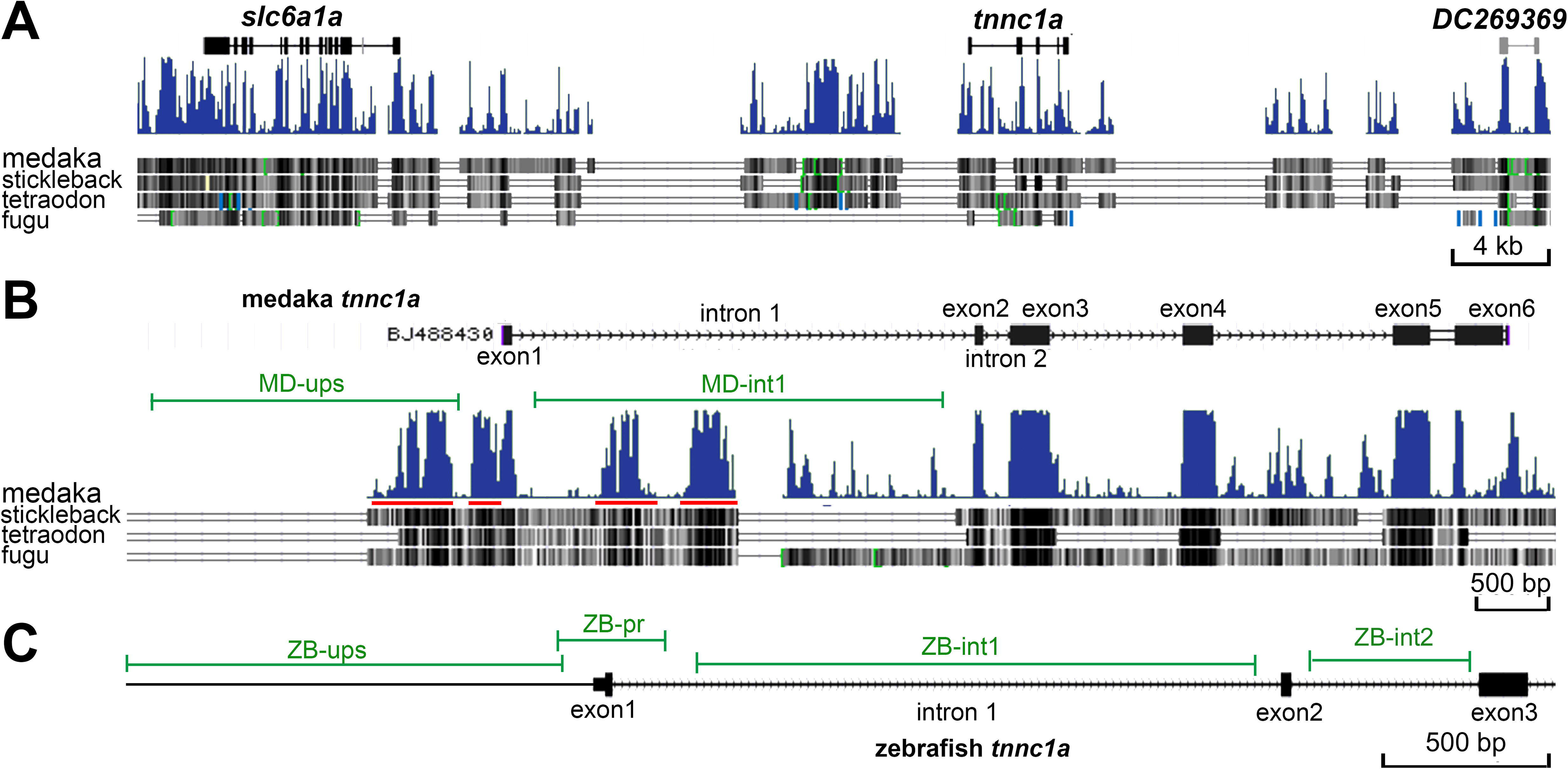
Conserved non-coding sequences in the teleost *tnnc1a* loci. **A.** The *tnnc1a* loci are flanked by neuronal-specific *slc6a1a* (Arcanjo et al., 2018) and EST DC269369 of unknown function expressed in fin regeneration (Katogi et al., 2004). Histograms show sequence conservation—the higher peaks, the better conservation among five teleost species (medaka, stickleback, tetraodon, fugu, and zebrafish). **B.** High magnification shows the exon-intron structures of the *tnnc1a* loci. The green lines indicate the enhancer candidate-containing upstream (ups) and intron 1 (int1) regions that were tested in this study, with red lines indicating the highly conserved non-coding regions. **C.** A local map of the 5’ end of the zebrafish *tnnc1a* gene. Green lines indicate the cis-regulatory element-containing upstream (ZB-ups), promoter (ZB-pr), and intronic (ZB-int1, ZB-int2) regions tested in this study.

Nevertheless, the cis-regulatory elements of zebrafish *tnnc1a* may still be within the corresponding upstream and intronic regions like in other fish, as in the case of the *ponli* enhancer (Fang et al., 2017).

Tissue-specific transcription is often mediated by enhancers, which can localize either upstream or downstream of the transcription start sites in distance. By contrast, basal promoters, normally located within 100 bp from transcription start sites, do not confer tissue specificity (Lenhard et al., 2012). Thus, the enhancer candidates of *tnnc1a* most likely reside in the conserved non-coding sequences that are 100 bp away from the transcription start site. To determine whether any of these distant conserved non-coding sequences (Fig. 3B, red lines) confer cardiac transcription, we first evaluated large stretches of DNA that contain conserved sequences before narrowing down the enhancer regions by truncation analysis. So, we assessed the transcriptional activities of a 1,150-bp medaka upstream DNA, a 1,649-bp medaka intron 1 DNA, a 1,299-bp zebrafish upstream DNA, and a 1,658-bp zebrafish intron 1

DNA (Fig. 3B and 3C) using an *in vivo* transgenic enhancer-reporting assay we developed previously (Fang et al., 2017) (Fig. 4A). Using this assay the transcriptional activities of a candidate enhancer can be inferred from the expression patterns of a GFP reporter gene (Fig. 4B).

**Figure 4.**
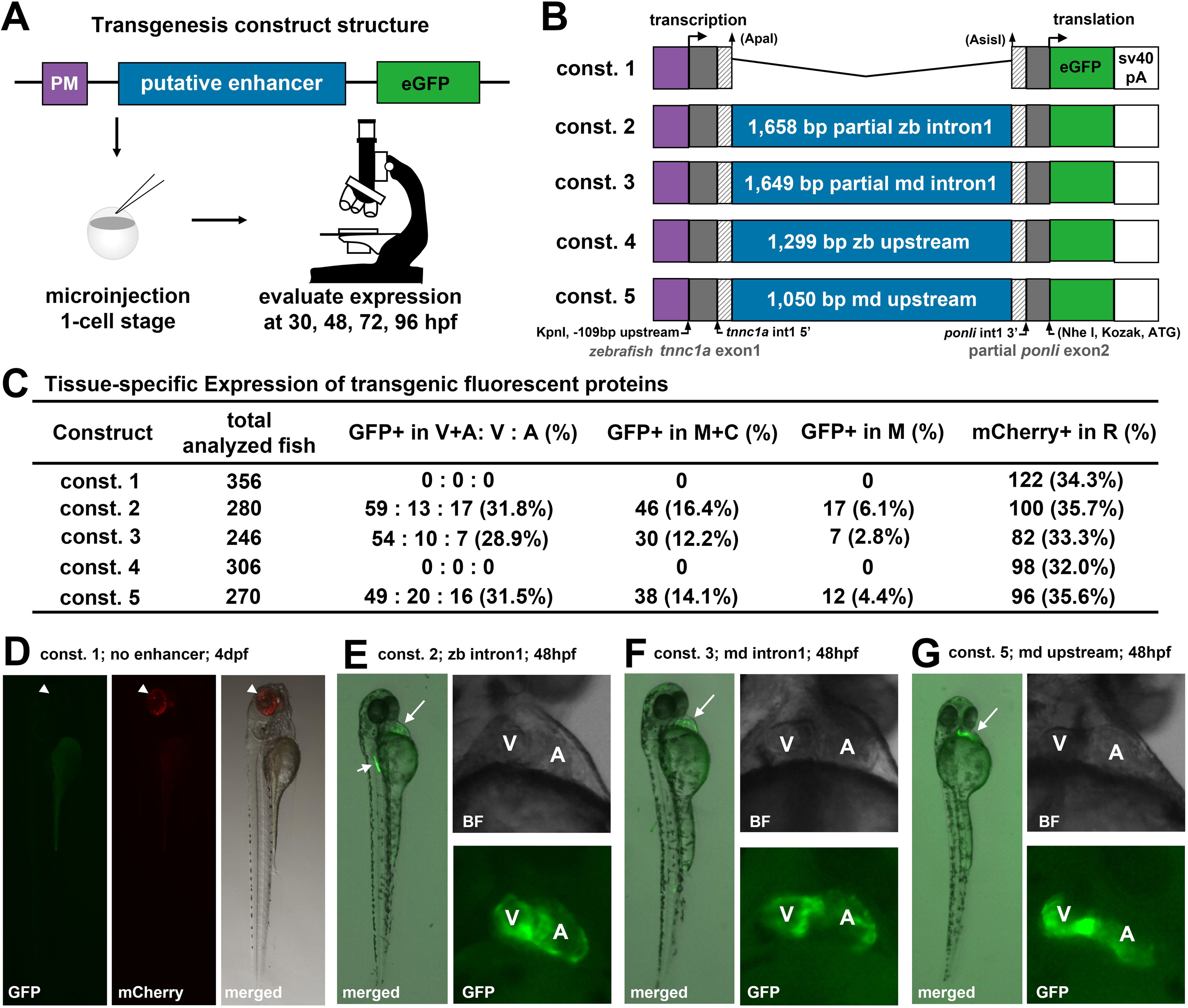
Identifying the cis-regulatory elements of zebrafish and medaka *tnnc1a* genes. **A.** The *in vivo* enhancer-testing assay. With this assay, an active enhancer can drive GFP expression in the corresponding tissue in 15-45% of injected embryos. PM, promoter; **B.** The structures of transgenic constructs 1-5 differ in enhancer candidates they contain, ranging from no enhancer, 1,658-bp zebrafish intron 1, 1,649-bp medaka intron 1, 1,299-bp zebrafish upstream DNA, and the 1,050-bp medaka upstream DNA, respectively. The 327-bp tnnc1a basal promoter region covers a 109-bp upstream DNA, exon 1, and 5’ end of intron 1. **C.** A table summarizes the expression patterns of constructs 1-5 and an mCherry-expressing coinjected control construct: GFP expression at 48 hpf, and mCherry at 4dpf. V, ventricle; A, atrium; M, muscles; R, retina. **D.** A whole body view of GFP and mCherry expression in a 4-dpf embryo injected with construct 1. **E-G.** The views of the whole body and the heart of 48-hpf embryos injected with construct 2 (E), construct 3 (F), and construct 5 (G) showed GFP expression in the heart. Muscle expression was occasionally expressed (E). A, atrium; V, ventricle.

By this assay, we found that the 1,658-bp zebrafish intron 1 DNA, the 1,150-bp medaka upstream DNA, and the 1,649-bp medaka intron 1 DNA displayed cardiac enhancer activities because GFP was expressed in the heart in about 30% of injected embryos.

This ratio is consistent with that of a co-injected mCherry-expressing control construct, which is driven by the green opsin gene promoter (Fig. 4). GFP expression emerged as early as 30 hpf (Fig. 4D) when the heart tube started to differentiate into two chambers. At 48 hpf, GFP was expressed in both heart chambers with stronger signals in the ventricle (Fig. 4), echoing the ventricular-biased mRNA expression of endogenous *tnnc1a* gene as revealed by in situ analysis (Fig. 2). Conversely, constructs that lack a testing enhancer or contain the 1,299-bp zebrafish upstream

DNA did not display any transcriptional activity (Fig. 4C), even though about 30% of embryos expressed mCherry in the retina as a control. These results suggest that the zebrafish *tnnc1a* basal promoter region by itself is not sufficient to activate cardiac transcription; however, the combination of the *tnnc1a* basal promoter with zebrafish *tnnc1a* intron 1 DNA, medaka *tnnc1a* upstream DNA, or medaka *tnnc1a* intron 1 DNA was sufficient to recapitulate the cardiac expression of the *tnnc1a* gene.

To determine whether the 1,658-bp DNA of zebrafish intron1, 1,050-bp medaka upstream DNA, and 1,649-bp DNA of medaka *tnnc1a* intron 1 are sufficient to confer the cardiac transcriptional specificity as enhancers, we next generated and evaluated chimeric transgenes’ transcriptional activities by combining these enhancer candidates separately with the basal promoter region of the *ponli* gene, which is restrictively expressed in red, green, and blue (RGB) cone photoreceptors of the retina (Fang et al., 2017; Zou et al., 2010). As expected, the resulting chimeric transgenes (Fig. 5) were activated in both heart chambers in embryos starting at 72 hpf in about 30% of injected embryos. Occasionally, in 4% of injected embryos, retinal GFP expression was observed, but the retinal expression was not observed in all cardiac GFP-expressing embryos (Fig. 5. B-J). The cause of this sporadic retinal expression is unclear, but it can be reasoned that the ponli basal promoter also contains some latent retinal specific activity, which can be activated when the transgene is integrated into some rare permissive genomic locations (Fang et al., 2017). Nevertheless, no cardiac activity was observed when the ponli basal promoter was used alone. Of note, the cardiac GFP expression driven by the combination of the heterologous *ponli* basal promoter and *tnnc1a* enhancers did not start at 30 hpf (data not shown) as when the *tnnc1a* basal promoter was used as the basal promoter (Fig. 4). This delay of expression was likely because the TATA-box-less *ponli* promoter is weak (Fang et al., 2017). Together, these results suggest that, when combined with a heterologous basal promoter, the 1,658-bp DNA of zebrafish intron1, the 1,050-bp medaka upstream DNA, and the 1,649-bp DNA of medaka *tnnc1a* intron 1 all harbor enhancers that can sufficiently and specifically activate pan-cardiac transcription.

**Figure 5.**
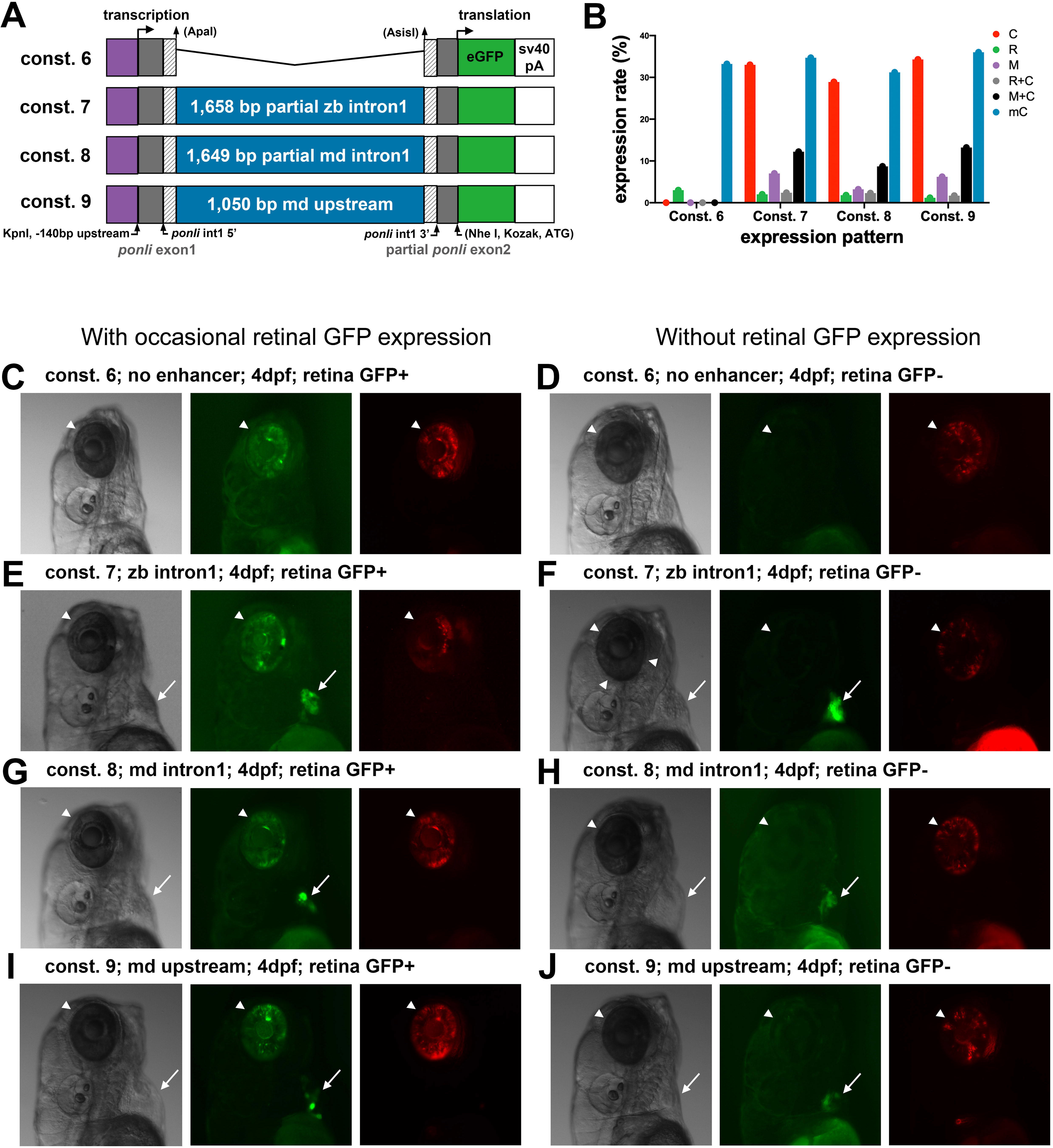
Identification of enhancer regions by chimeric transgene analysis. **A.** The transgenic constructs 6-9 are the combinations of the *ponli* basal promoter region with no enhancer, the 1,658-bp zebrafish intronic 1 DNA, the 1,649-bp medaka intron 1 DNA, and the 1,050-bp medaka upstream DNA to assess the transcriptional specificity of these enhancer candidates. **B.** Histograms illustrate GFP expression at 4 dpf in injected embryos (coinjected with the mCherry-expressing control construct) in the muscles (M), heart ventricle (V), heart atrium (A), and retina (R). **C-J.** Representative expression patterns at 4 dpf of embryos injected with constructs 6-9, either with or without sporadic retinal expression of GFP observed in 4% of injected fish. Arrowheads indicate retinal expression; arrows indicate cardiac expression. mCherry expression was from a co-injected control construct driven by the RH2-1 green opsin cis-regulatory elements.

### Narrowing down the *tnnc1a* enhancer regions

Enhancers can be either short, with TF binding sites packed densely, or long, with TF binding sites scattered more broadly (Yanez-Cuna et al., 2013). To further define the three regions that harbor the *tnnc1a* enhancers in medaka and zebrafish, we next truncated these enhancer regions and tested the transcriptional activities of the resulting constructs. We found that a 509-bp mid region of zebrafish intron 1, a 668-bp of medaka intron 1, and 241-bp medaka upstream DNA could recapitulate the cardiac expression (Fig. 6. A-F). Two stable transgenic fish lines were also obtained—*Tg(tnnc1a^medaka ups-1,050 bp^:GFP)^pt188^* and *Tg(tnnc1a^zebrafish int1-509 bp^:GFP)pt189*, which show cardiac expression only without muscle expression, suggesting that sporadic muscle expression in injected founder fish is nonspecific entopic expression (Fig. 6. G, H). These results suggested that the tnnc1a enhancers are harbored in these narrower regions.

**Figure 6.**
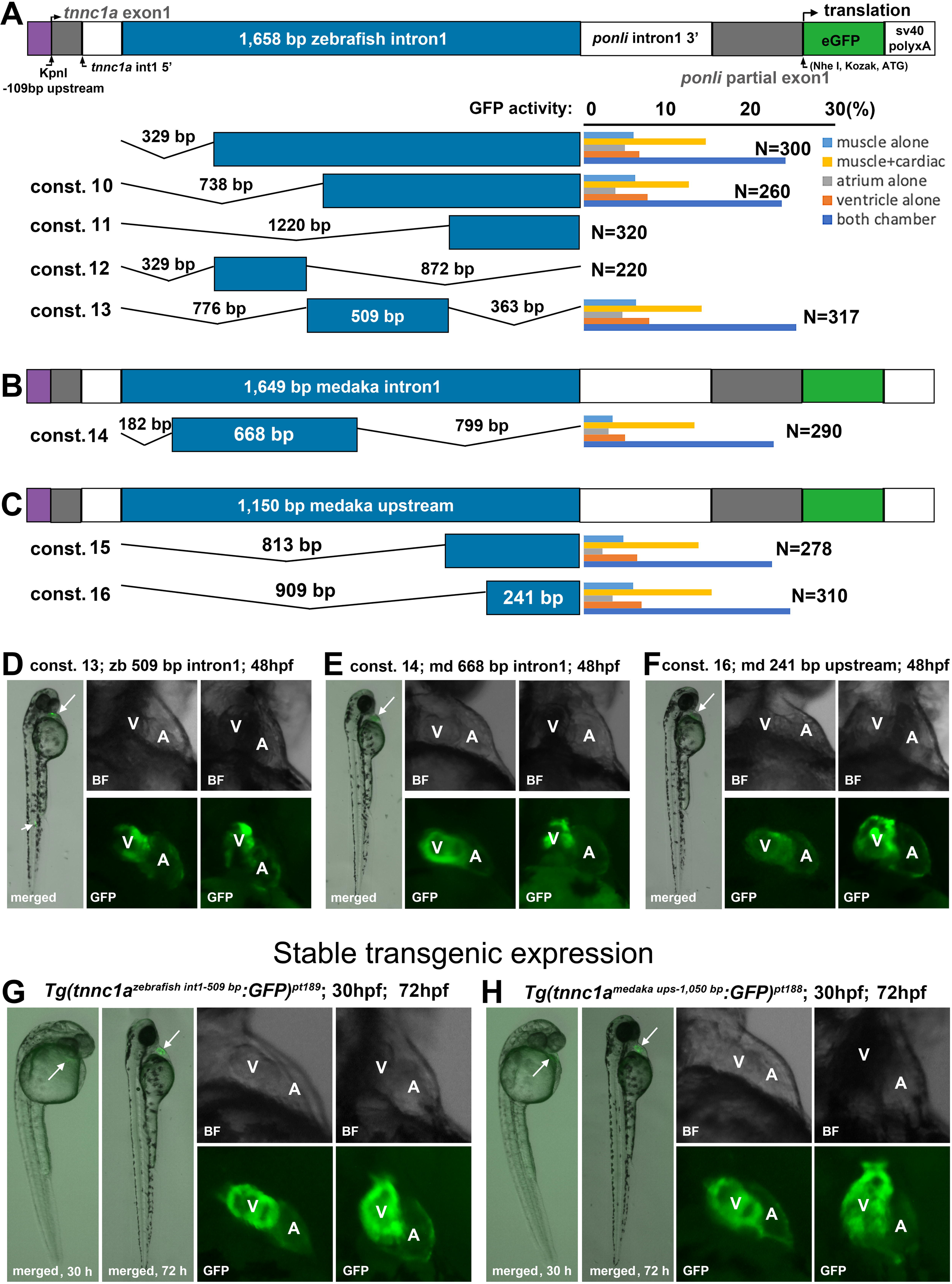
Narrowing down the regions that harbor the zebrafish and medaka *tnnc1a* enhancers**. A-C.** Schematics illustrate that deletion analysis narrowed down the enhancer-harboring regions (blue boxes) from the initial broader regions: 1,658-bp zebrafish intron 1 (A), 1,549-bp medaka intron 1 (B), and 1,150-bp medaka upstream DNA (C). Histograms represent the percentage of GFP-positive embryos in injected fish, quantified at 48 hpf per tissue categories: V, ventricle; A, atrium; M, muscles; M+C, both muscle and cardiac. **D-F.** GFP expression at 48 hpf in the heart of an embryo injected with constructs 13, 14, and 16. **G-H.** GFP expression in fish of stable transgenic lines *Tg(tnnc1a^medaka^ ^ups-1,050^ ^bp^:GFP)^pt188^* and *Tg(tnnc1a^zebrafish^ ^int1-509 bp:^GFP)^pt189^*.

### Conserved sequence motifs in *tnnc1a* basal promoters and enhancers

As cis-regulatory elements, *tnnc1a*’s basal promoters and enhancers regulate its transcription by recruiting transcription factors (TFs) via short sequence motifs. Identification of these motifs will provide handles to study what trans-regulatory TFs interact with cis-regulatory elements to control *tnnc1a* transcription. To identify these sequence motifs, we compared the sequences of the promoter and enhancer regions of teleost *tnnc1a* genes using MEME and Improbizer. In the promoter regions of the *tnnc1a* genes (arbitrarily defined as a region from -109 bp to 160 bp downstream of the 5’ splicing site of *tnnc1a* intron 1, which overlap with the 60-bp *tnnc1a* basal promoter predicted in the Eukaryotic Promoter Database (https://epd.expasy.org/cgi-bin/epd/get_doc?db=drEpdNew&format=genome&entry=tnnc1a_1)(Dreos et al., 2017; Dreos et al., 2015), two motifs of non-coding sequence are highly conserved among all five fish species: a CG-rich site at - 46 bp to - 37 bp, and a TATA box at – 34 bp to – 20 bp (Fig. 7). As predicted by JASPAR and Transfac, these sites are targets of general transcription factors SP1 and TBP (TATA-binding protein) of TFIID, respectively. These sites are often associated with basal promoters (Landolin et al., 2010). SP1 can recruit TBP to assemble the RNA polymerase II preinitiation complex (Bhuiyan and Timmers, 2019; Butler and Kadonaga, 2002).

**Figure 7.**
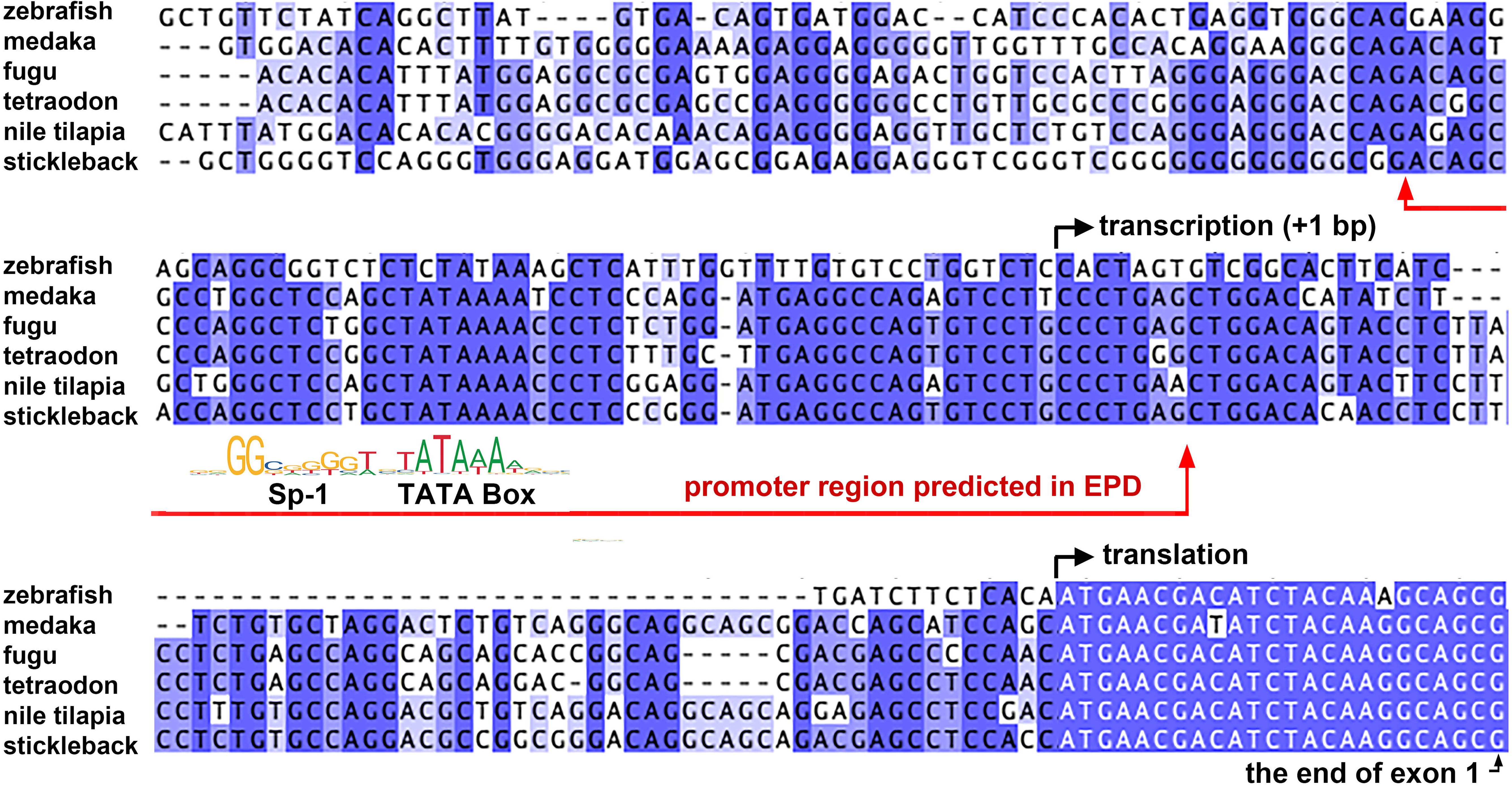
Sequence conservation in the basal promoters of the *tnnc1a* genes. **A.** Alignment of sequences at the promoter regions of six teleost fish revealed conserved motifs for SP-1 and TBP. The red line indicates the promoter region predicted in the EPD (Eukaryotic promoter database).

Since basal promoters normally span a short region (50–80 bp) around the transcription start site, we propose that the basal promoters of *tnnc1a* likely span the regions from about -60 to about +50, a region consistent with the basal promoter predicted in the Eukaryotic Promoter Database. The presence of binding sites for general transcription factors SP1 and TBP is consistent with the expectation that *tnnc1a* basal promoters do not play a role in regulating cardiac transcriptional specificity.

Similarly, short conserved sequence motifs are present in the medaka and zebrafish *tnnc1a* upstream and intronic enhancer regions (Fig. 8A, B). These conserved motifs match the consensus sequences of many TFs, as predicted by Transfac, TOMTOM, and JASPAR. Among these TFs are known cardiac TFs, such as GATA4, MEF2C, MYOD1, and Nkx2-5. The motif conservation between medaka’s upstream and zebrafish’s intronic enhancers suggests that the upstream and intronic enhancers likely share ancestry during evolution.

**Figure 8.**
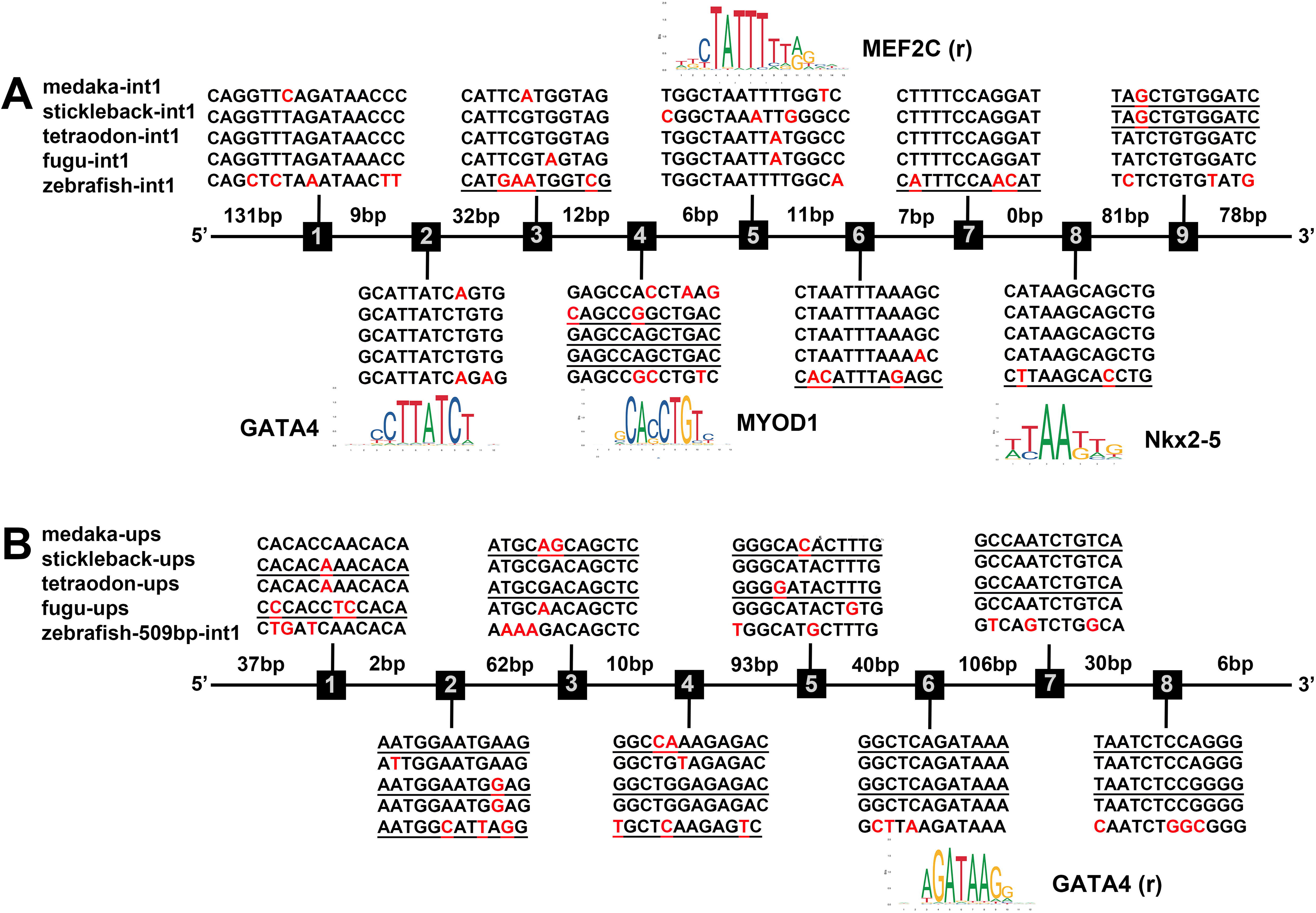
A. A schematic showing that the 509-bp zebrafish *tnnc1a* intronic enhancer contains nine consensus motifs conserved with other teleost fish (medaka, stickleback, tetraodon, and fugu) *tnnc1a* intron1, whose sequence alignments are presented either above or below the schematic. The red letters represent unconserved nucleotides. Underlines indicate a reversed orientation relative to the motif orientation of species. Conserved motifs are potential targets of many transcription factors as identified by JASPAR. The consensus motif sequence of known transcription factors (bottom) and the *tnnc1a* enhancer motifs (top) are aligned to reveal the potential binding sites. **B.** Comparison between the upstream and intronic enhancers. The intervals between motifs are based on medaka sequences.

## Discussion

In this study, we identified and characterized the enhancers of the cardiac-specific *tnnc1a* genes of medaka and zebrafish. By comparing these *tnnc1a* enhancers with the enhancers of other cardiac genes, we also revealed common enhancer motifs that play roles in cardiac-specific transcription, providing insights into the general principles governing pan-cardiac transcriptional regulation.

### Teleost *tnnc1a* enhancers and basal promoters

The sizes of zebrafish, medaka, stickleback, fugu, and tetraodon genomes vary significantly, ranging from 1.4 gigabases for zebrafish as the largest to 350 megabases for tetraodon as the smallest (Aparicio et al., 2002; Howe et al., 2013; Jaillon et al., 2004; Kasahara et al., 2007; Peichel et al., 2017). Despite these variations, all five fish species contain an enhancer in the first introns of the *tnnc1a* genes. Interestingly, an additional upstream *tnnc1a* enhancer is present in medaka (likely also in stickleback, fugu, and tetraodon per sequence conservation), but not found within 1,300 bp upstream of the zebrafish tnnc1a gene (we do not rule out the possibility that another enhancer is present in a more distant region, which was not tested in this study). The upstream medaka *tnnc1a* enhancer appears equivalent to the intronic enhancer because both can specifically drive pan-cardiac expression in zebrafish, whether combined with the cognate *tnnc1a* basal promoter or the heterologous *ponli* basal promoter. It is unclear whether the upstream and intronic enhancers regulate cardiac expression together in a synergistic fashion.

The cardiac specificity of transcription is likely solely determined by *tnnc1a*’s enhancers, with no contribution from the teleost *tnnc1a* basal promoters, which contain a binding site for general transcription factor SP1 site and a TATA box for TBP. In addition, the broader basal promoter region used in constructs, i.e., from -109 bp to 160 bp downstream of the first intron’s 5’ splicing site, does not appear to contain additional sequence motifs that may contribute to the cardiac specificity of *tnnc1a* expression because it alone could not drive cardiac expression. Having a TATA box to better recruit TFIID via TBP and an SP1 site to facilitate the recruitment of TBP, the *tnnc1a* basal promoters are supposed to be a stronger basal promoter than the TATA-less *ponli* basal promoter; this may explain why GFP was not observed in early heart tube (30 hpf) when the GFP reporter gene was driven by the combination of *tnnc1a* enhancers and the *ponli* basal promoter.

### Enhancer regulation of mammalian *tnnc1* and teleost *tnnc1a* and *tnnc1b*

Unlike the cardiac-specific expression of teleost *tnnc1a*, the mammalian counterpart *tnnc1* is expressed in both cardiac muscles and slow-twitch skeletal muscles. The cardiac expression is driven by its upstream enhancer (Parmacek et al., 1992), whereas its expression in slow-twitch skeletal muscles is driven by an intronic enhancer (Parmacek et al., 1994). This suggests that ancestral *tnnc1* genes were expressed in both cardiac and slow-twitch skeletal muscles, driven by independent enhancers. However, in teleost, due to the teleost-specific whole genome duplication that is expected to contribute to teleost complexity (Glasauer and Neuhauss, 2014), the cardiac and slow-twitching functions became subfunctionalized in the resulting *tnnc1a* and *tnnc1b* genes, with *tnnc1a* restrictively expressed in the heart and *tnnc1b* in slow-twitching skeletal muscles in embryos (Sogah et al., 2010) and in both heart and muscles in adult zebrafish (Genge et al., 2013). To achieve the cardiac restrictive expression of *tnnc1a*, the skeletal enhancers were likely converted into a redundant cardiac enhancer in medaka (likely also in stickleback, fugu, and tetraodon) or inactivated in zebrafish. Along the same line of thinking, it can be speculated that the original cardiac enhancer of *tnnc1b* became inactive at least in the embryonic heart during evolution. Such differential expression regulation of tnnc1a and tnnc1b may be beneficial because fish, as ectotherm animals, the heart and skeletal muscles may require different regulation by tnnc1 to maintain different physiological properties (Genge et al., 2013; Genge et al., 2016; Gillis et al., 2007; Klaiman et al., 2011).

## Acknowledgments

This work was supported by NIH grant EY025638 to XW and NIH core grant P30EY008098 to the Department of Ophthalmology. XW was also supported by the Department of Ophthalmology at the University of Pittsburgh, the Eye and Ear Foundation of Pittsburgh, and an unrestricted grant from Research to Prevent Blindness to the Department of Ophthalmology at the University of Pittsburgh. JB was a graduate student of the joint training program of the Xiangya Medical School and the University of Pittsburgh, and she was financially supported by Xiangya Medical School.

